# Micromagnetic Stimulation (μMS) Controls Dopamine Release: An *in vivo* Study Using WINCS *Harmoni*

**DOI:** 10.1101/2023.05.25.542334

**Authors:** Renata Saha, Abhinav Goyal, Jason Yuen, Yoonbae Oh, Robert P. Bloom, Onri J. Benally, Kai Wu, Theoden I. Netoff, Walter C. Low, Kevin E. Bennet, Kendall H. Lee, Hojin Shin, Jian-Ping Wang

**Affiliations:** Department of Electrical and Computer Engineering, University of Minnesota, Minneapolis, MN, United States; Department of Neurologic Surgery, Mayo Clinic, Rochester, MN, United States; Medical Scientist Training Program, Mayo Clinic, Rochester, MN, United States; Deakin University, IMPACT – the Institute for Mental and Physical Health and Clinical Translation, School of Medicine, Barwon Health, Geelong VIC 3216, Australia; Department of Biomedical Engineering, Mayo Clinic, Rochester, MN, United States; Department of Biomedical Engineering, University of Minnesota, Minneapolis, MN, United States; Department of Neurosurgery, University of Minnesota, Minneapolis, MN, United States; Division of Engineering, Mayo Clinic, Rochester, MN, United States

**Keywords:** neurotransmitters, μcoils, micromagetic stimulation, dopamine, MFB, striatum, FSCV, WINCS Harmoni

## Abstract

**Objective:** Research into the role of neurotransmitters in regulating normal and pathologic brain functions has made significant progress. Yet, clinical trials that aim to improve therapeutic interventions do not take advantage of the *in vivo* changes in the neurochemistry that occur in real time during disease progression, drug interactions or response to pharmacological, cognitive, behavioral, and neuromodulation therapies. In this work, we used the WINCS *Harmoni* tool to study the real time *in vivo* changes in dopamine release in rodent brains for the micromagnetic neuromodulation therapy.

**Approach:** Although still in its infancy, micromagnetic stimulation (μMS) using micro-meter sized coils or microcoils (μcoils) has shown incredible promise in spatially selective, galvanic contact free and highly focal neuromodulation. These μcoils are powered by a time-varying current which generates a magnetic field. As per Faraday’s Laws of Electromagnetic Induction, this magnetic field induces an electric field in a conducting medium (here, the brain tissues). We used a solenoidal-shaped μcoil to stimulate the medial forebrain bundle (MFB) of the rodent brain *in vivo*. The evoked *in vivo* dopamine releases in the striatum were tracked in real time by carbon fiber microelectrodes (CFM) using fast scan cyclic voltammetry (FSCV).

**Results:** Our experiments report that μcoils can successfully activate the MFB in rodent brains, triggering dopamine release *in vivo*. We further show that the successful release of dopamine upon micromagnetic stimulation is dependent on the orientation of the μcoil. Furthermore, varied intensities of μMS can control the concentration of dopamine releases in the striatum.

**Significance:** This work helps us better understand the brain and its conditions arising from a new therapeutic intervention, like μMS, at the level of neurotransmitter release. Despite its early stage, this study potentially paves the path for μMS to enter the clinical world as a precisely controlled and optimized neuromodulation therapy.

## 1. Introduction

Chemical signalling mediated by an intricate balance of specific neurotransmitters is necessary for normal functioning of the brain. Diseases, injury or new therapeutic interventions could disrupt this balance of neurotransmitters and the results can be devastating. For instance, low dopamine levels are linked to the following medical conditions – anxiety, depression, tremors, fatigue, and cognitive imbalance [1–6]. In a similar context, low levels of serotonin are associated with insomnia, migraine headaches, suicidal ideation among other conditions[7–9]. These and other brain disorders associated with neurotransmitter imbalance affect one billion people globally [10]. Therefore, accurate measurement of these neurotransmitter concentrations in patients during onset of diseases, recovering from injuries or testing new therapies is of utmost importance [11].

The microdialysis is widely used for measuring neurochemical concentrations with high selectivity and specificity [12,13]. However, this technique has drawbacks when compared to electrochemical methods such as fast scan cyclic voltammetry (FSCV) in terms of its large microdialysis probe size (> 200 μm) and limited temporal resolution (< 1 min) [14–16]. Electrochemical techniques such as FSCV are well suited for measuring extracellular dopamine concentrations in real time with much better spatiotemporal resolutions (∼milliseconds) and detection sensitivity (< 5 nM). The carbon fiber microelectrode (CFM) probe used in FSCV is of much smaller diameter (< 10 μm), causing less damage to brain tissues compared to microdialysis probes. The voltage applied to the CFM is initially kept at a fixed resting potential and then gradually ramped up to an electric potential necessary to oxidize and reduce electroactive species (e.g. dopamine) before the potential is returned to the resting potential again [17]. This voltage sweep is fast (< 10 ms) and is repeated every 100 ms (equivalent to a scanning frequency of 10 Hz). The voltammogram provides a chemical signature which is unique to the electroactive species (dopamine, serotonin, acetylcholine etc.) and can be used to study their phasic changes in extracellular concentrations.

Neuronal dopamine release from the mammalian midbrain can be distinctly characterized into 2 patterns: tonic activity and phasic burst activity [18]. Tonic activity is observed when extrasynaptic dopamine is released spontaneously by pacemaker-like firing of midbrain neurons into the striatum. This modulates behavioral flexibility by maintaining a relatively homeostatic extracellular concentration of dopamine. Whereas during phasic burst activity, transient dopamine is released at the synapse, triggering neural plasticity. However, FSCV can only detect phasic changes of dopamine or neurotransmitter concentrations, as large capactitive currents at the CFM surface need to be subtracted to resolve the Faradaic current for precise voltammetric measurements [18].

Micromagnetic neurostimulation was first experimentally implemented in an *ex vivo* study on rabbit retinal neurons using commercially available solenoidal-shaped microcoils (μcoils) [19]. Since then, attempts have been made by several groups to use similar μcoil prototypes in several experimental settings, both *in vitro* and *in vivo*, to showcase the efficacy of micromagnetic neurostimulation [20–25]. Furthermore, different designs of μcoils, solenoidal [20], planar [22,26], V-shaped [27] or trapezoidal [27], with or without soft magnetic material cores [21], have been investigated. In this neuromodulation technique, μcoils are the devices that are powered by time-varying currents at varying amplitudes, pulse widths (PW) and frequencies. According to Faraday’s Laws of Electromagnetic Induction, these current carrying μcoils generate a magnetic field, which in turn induces an electric field on a conducting surface (here, biological conductors such as neural tissues.). Since the induced electric field that activates the neural tissues are not in direct galvanic contact with the tissues, these μcoils exhibit less implant surface corrosion due to biofouling [21]. The induced electric field is highly directional in nature causing spatially-selective activation of the neural tissue from these μcoils [20]. Furthermore, in a recent numerical study, these μcoils have shown significantly less radio frequency (RF) heating (antenna effect) from a 1.5 Tesla magnetic Resonance Imaging (1.5T-MRI) environment compared to traditional deep brain stimulation (DBS) leads [28]. Therefore, these μcoils are MRI-safe, enabling patients with these implants to undergo an MRI scan after insertion of these μcoil implants in their brains.

Micromagnetic neurostimulation is quite recent in its origin and is still in its infancy. Tremendous efforts are yet to be made to develop the μcoils to be fit for testing on non-human primates to gradually progress them into clinical trials. Being a relatively new neuromodulation therapy, a study on the associated neurotransmitter releases is required. In this work, we made an attempt to study how micromagnetic stimulation of the substantia nigra-striatal nerve fibers in the medial forebrain bundle (MFB) in a rodent brain releases dopamine *in vivo*. Upon stimulation from the μcoils, we quantified the phasic dopamine responses in real time from the striatum using FSCV. This work is a proof-of-concept study on rodent brains *in vivo*, demonstrating the feasibility of μcoils in activating the MFB fibers and the neuronal dopaminergic pathway thereby quantifying the dopamine release from the striatum.

## 2. Materials and Methods

### 2.1. The μcoil Probe Fabrication

In this *in vivo* study, we used the Magnetic Pen (MagPen) as the μcoil implant (μcoil dimension = 1 mm × 0.6 mm × 0.5 mm) [20,21]. The commercially available solenoidal μcoil of model no. TDK Corporation MLG1005SR10JTD25 was soldered at the tip of the green printed circuit board (PCB) of length 3 cm. To minimize brain tissue damage during implantation of the MagPen probe, the thickness of the PCB was reduced to 0.4 mm. To determine if the MagPen orientation with respect to the MFB fibers in the rodent brain affected dopamine release from the striatum, we fabricated the prototype in two orientations: Type <u>H</u>orizontal (Type H or MagPen-Type H) and Type <u>V</u>ertical (Type V or MagPen-Type V) (see Fig. S1(a)). The width of the μcoil tips for MagPen-Type H and MagPen-Type V were optimized to be at 1.7 mm and 1.4 mm respectively. The complete image of the MagPen prototype has been reported in Fig. S1(b) (see Supplementary Information S1). To ensure complete biocompatibility in addition to an antileakage current and waterproof sealing for the probe, the tip of the MagPen was encapsulated in a 10 μm thick coating of Parylene-C (SCS-Labcoater Parylene-C deposition system). Successful coating for the MagPen prototype was determined once the impedance between the MagPen and a salt bath was > 5MΩ [20,21].

### 2.2. The Electrical Circuit Characterization of the μcoil

Resistance (R), inductance (L) and capacitance (C) values of the μcoil implant were measured using an LCR meter (Model no. BK Precision 889B; see Table S1 in Supplementary Information S1) at two frequencies (120 Hz and 1 kHz). The frequency of operation for the μcoil in this work was around 60-130 Hz compared to other reported works with MagPen where the frequency of operation for the μcoil was at 1-5 kHz [20,21]. The frequency of operation around 60-130 Hz contributed to the benefit for micromagnetic operation on MFB fibers in the rodent brain as compared to the LCR meter measurements at 1 kHz. The DC resistance (R_DC_), the series inductance (L_s_) and the parallel capacitance (C_p_) values remained almost the same for both frequency measurements (120 Hz and 1 kHz; see Table S1). The series capacitance (C_s_) value increased, and the parallel inductance (L_p_) value decreased by several orders (see Table S1) at 120 Hz. Therefore, reactance from 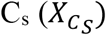 offered lower resistance to the current flow in the series RL branch and L_p_ offered higher resistance to the parallel LC branch 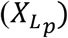 resulting in complete flow of current through the series RL branch of the μcoil (see Fig. S1(c)). Therefore, the μcoil in MagPen acted as a series RL circuit (see Fig. S1(c)).

The induced electric field in micromagnetic activation of neurons is a function of L_s_. The electromotive force (emf or *v(t)*) induced in the neural tissues is expressed as: 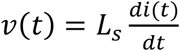. This emf is generated due to a time-varying magnetic field (***B(t)***) as per Faraday’s Laws of Electromagnetic Induction and 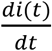 is the time derivative of the time-varying current (***i(t)***) through the inductor or μcoil. This has been used to study the effect of micromagnetic neurostimulation on the MFB fibers of the rat brain to trigger dopamine release in rats (see section 3.2). The resistance (R) contributes to the Joule heating on the neural tissues from the μcoil (*heat dissipated = i*(*t*)^2^*R*_*DC*_*t*). Our numerical calculations and independent experiments corroborate with existing literature which report that the temperature increase from these μcoils on neural tissues upon micromagnetic stimulation remains below 1°C [21,22,25].

### 2.3. Fabrication of Carbon Fiber Microelectrode (CFM)

Carbon fiber microelectrodes (CFMs) (see Fig. 1(a)-i) were fabricated using a standardized design at Mayo Clinic, Rochester, MN [29,30]. The microelectrode involved isolating and inserting a single carbon fiber (AS4, diameter = 7 μm; Hexcel, Dublin, CA) into a silica tube (inner diameter (ID) = 20 μM, outer diameter (OD) = 90 μM OD, 10 μM thick coating with polyimide; Polymicro Technologies, Phoenix, AZ). The connection between the carbon fibers extending out of the silica tubing were sealed with epoxy resin. The carbon fiber in the silica tubing was then connected to a conductive wire composed of nitinol (Nitinol #1, an alloy of nickel and titanium; Fort Wayne Metals, IN) coated with a silver-based conductive paste [29]. The carbon fiber attached nitinol wire was then insulated with a polyimide tube (0.0089” ID, 0.0134” OD, 0.00225” WT; Vention Medical, Salem, NH) up to the carbon fiber sensing tip. The exposed carbon fiber was then trimmed under a dissecting microscope to a length of ∼50 μm. Teflon-coated silver (Ag) wire (A-M systems, Inc., Sequim, WA) was prepared as an Ag/AgCl counter-reference electrode by chlorinating the exposed tip in saline with a 9 V dry cell battery. CFM was pretested *in vitro* in a TRIS buffer (15 mM tris, 3.25 mM KCl, 140 mM NaCl, 1.2 mM CaCl_2_, 1.25 mM NaH_2_PO_4_, 1.2 mM MgCl_2_, and 2.0 mM Na_2_SO_4_, with the pH adjusted to 7.4) [31] prior to implantation.

**Figure 1.**
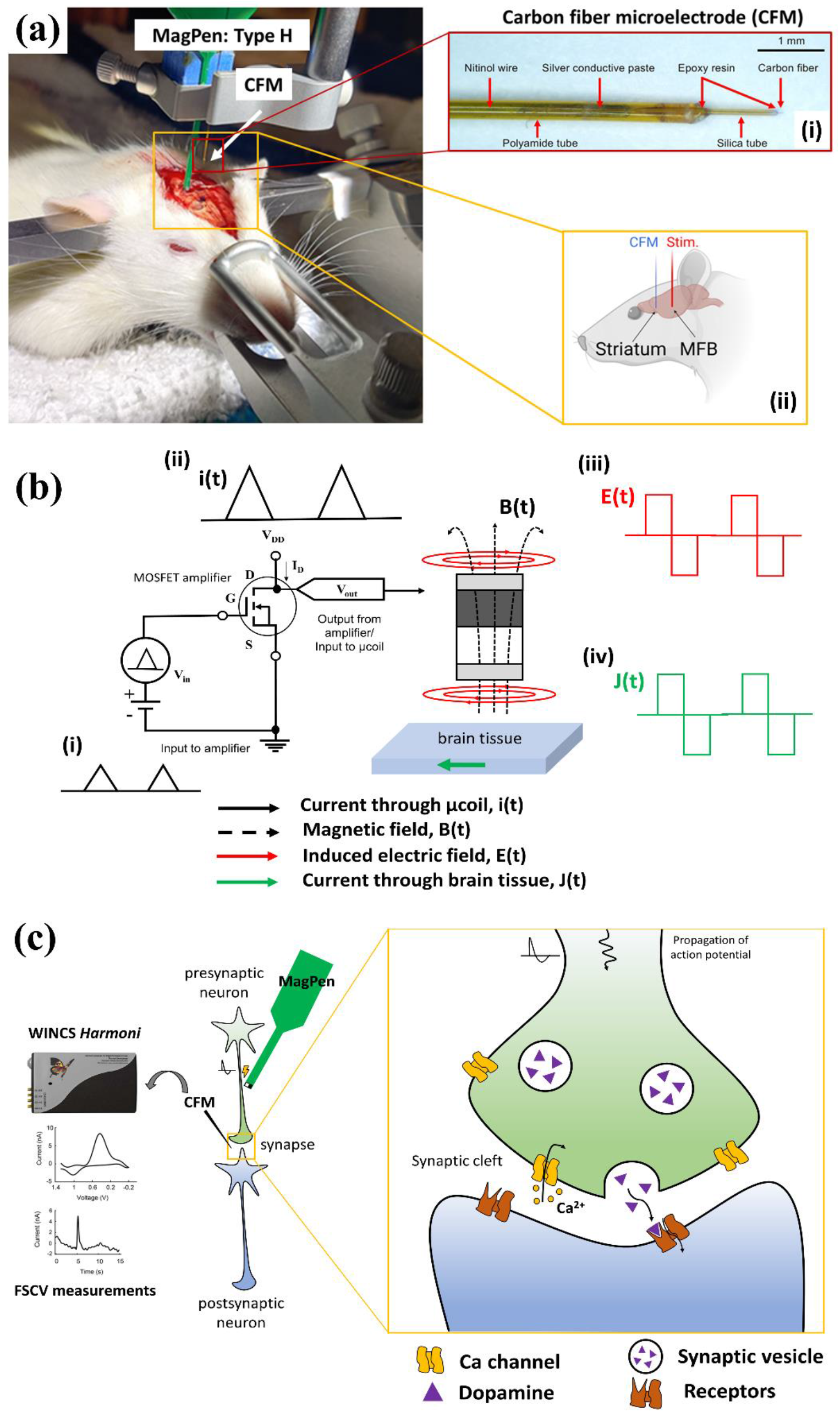
(a) *in vivo* experimental set-up where the MagPen is inserted into the medial forebrain bundle (MFB) and the carbon fiber microelectrode (CFM) is inserted into the striatum. (i) The zoom-in image of the CFM with its different parts labelled. (ii) Schematic overview of the regions of the brain being stimulated and recorded. Partially created with BioRender.com. (b) (i-iv) The signal flowchart through which the MFB is being stimulated by the MagPen-Type H. (c) Schematic overview of MagPen stimulation and CFM recording of dopamine at the synaptic cleft. Upon MagPen stimulation of the MFB, dopamine release is evoked in the dorsal striatum. This release is measured by the CFM linked with WINCS *Harmoni*.

### 2.4. Implantation of CFM and MagPen *in vivo*

The rat (Sprague Dawley, male, 250g, Envigo, United States) was anesthetized with urethane (1.5 g/kg i.p.; Sigma-Aldrich, St Louis, MO, USA) and administered buprenorphine (0.05-0.1 mg/kg s.c., Par Pharmaceutical, Chestnut Ridge, NY, USA) for analgesia. Upon complete anesthesia, rats were placed in a stereotaxic frame (David Kopf Instruments, Tujunga, CA, USA) with its body resting on a heating pad (41°C). Respiratory rate (RespiRAT, Intuitive Measurement Systems), hind-paw and tail pinch were used to monitor the physiological state and depth of anesthesia over the entire course of the experiment. Using a standard rat atlas [32], three trephine holes were drilled, the first for placement of a CFM into the striatum (AP 1.2 mm from bregma, ML 2.0 mm, and DV -5.1 mm from dura) (see Fig. 1(a)-i), the MagPen into the medial forebrain bundle (MFB) (AP -4.6 mm from bregma, ML 1.3 mm, and DV -9 mm from dura) (see Fig. 1(a)-ii), and a third for an Ag/AgCl reference/counter electrode into the contralateral cortex [33]. The animal study was reviewed and approved by the Institutional Animal Care and Use Committee (IACUC), Mayo Clinic, Rochester, MN. Protocol No.: A00003894.

### 2.5. The MagPen Driving Circuitry

The MagPen driving circuitry consists of a function generator (Model no. RIGOL DG1022Z) which is used to generate bursts of triangular waveforms of varied amplitudes and pulse widths (PW). The signal from the function generator is amplified by a Class-D amplifier (Model no. Pyramid PB717X) set at a constant gain A (see Fig. 1(b)-ii). The output from the amplifier powers the μcoil in the MagPen which is used to deliver micromagnetic stimulation. Using the trigger function in the function generator, the total stimulation time span and the wait time between each successive stimulation trial was controlled.

### 2.6. WINCS *Harmoni* for *in vivo* Neurotransmitter Recording

WINCS *Harmoni* [11] is a closed-loop neurochemical monitoring system that can monitor real-time changes in neurotransmitter levels upon *in vivo* stimulation. This research platform is a very convenient research tool that helps track changes in the brain process during disease progression and its response to pharmacologic, cognitive, behavioral and neuromodulation therapies by directly studying the brain chemistry.

### 2.7. Finite Element Modeling (FEM) of the Neural Implants

The bipolar twisted electrodes for *in vivo* electrical stimulation (Model no. Plastics One, MS 303/2, Roanoke, VA, USA) were modeled on ANSYS-Maxwell, electrostatic solver to compare the electric field contour plots between electrical and micromagnetic stimulation. The twisted electrode dimensions, tissue slab parameters, boundary conditions and the high-resolution tetrahedral mesh size used are detailed in Table 1. FEM simulations on Ansys-Maxwell, eddy current solver [34] (ANSYS, Canonsburg, PA, United States) were used to study the magnetic field and the induced electric field from the μcoils. It solves an advanced form of the T-Ω formulation of the Maxwell’s equations [35]. The ceramic core μcoil dimensions, tissue slab parameters, boundary conditions and the high-resolution tetrahedral mesh size used are detailed in Table 2. All simulations were done using the Minnesota Supercomputing Institute (MSI) at the University of Minnesota (8 cores of Intel Haswell E5-2680v3 CPU, 64×8=512 GB RAM and 1 Nvidia Tesla K20 GPU). The induced electric field values were then exported to be analyzed using a customized code written in MATLAB (The Mathworks, Inc., Natick, MA, USA).

**Table 1.**
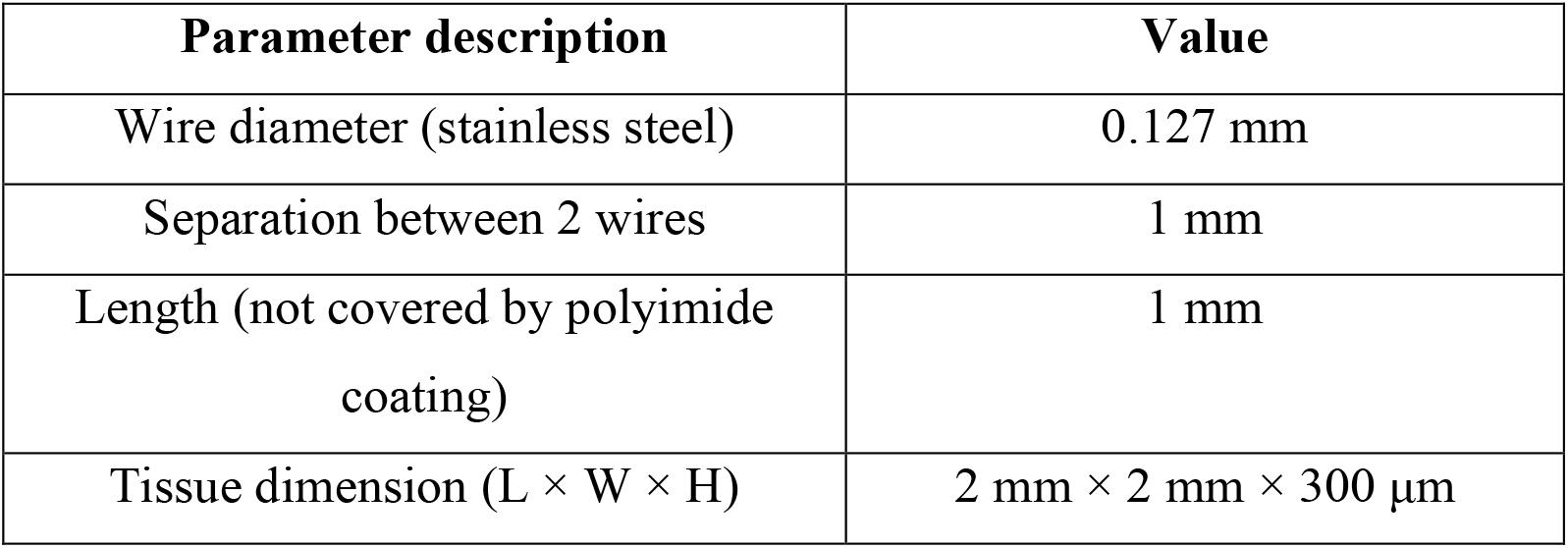
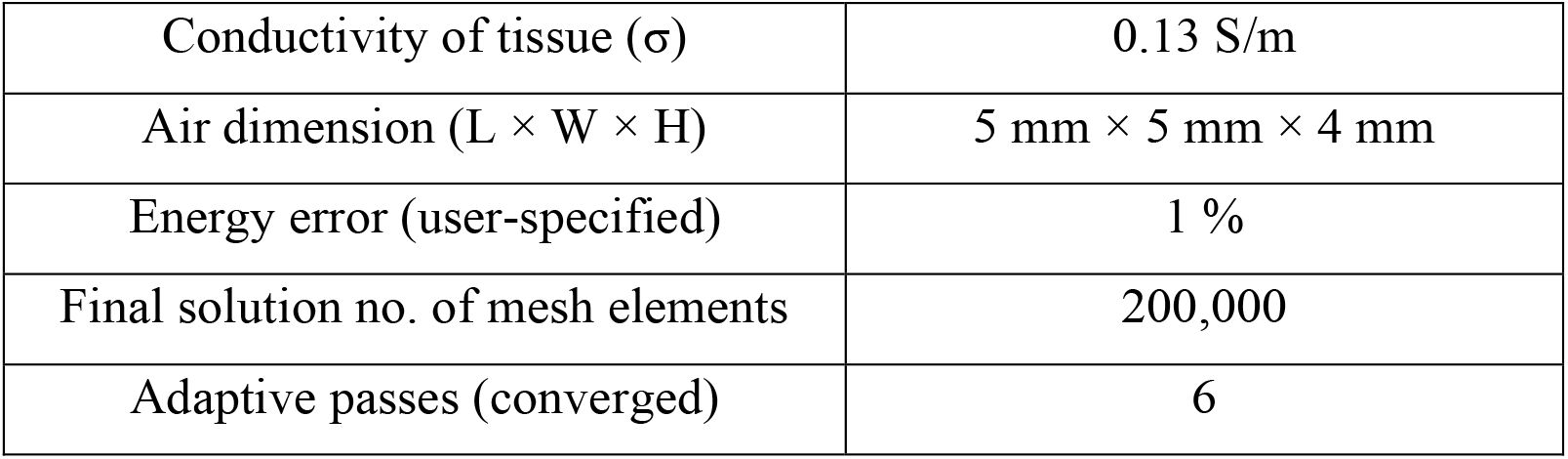
Modeling parameters for electrical electrode

**Table 2.** Modeling parameters for the μcoil

| Parameter description | Value |
| --- | --- |
| μcoil dimension ($L \times W \times H$) | 1 mm × 600 μm × 500 μm |
| no. of turns (N) | 21 |
| Wire diameter | 7 μm |
| Tissue dimension ($L \times W \times H$) | 4 mm × 4 mm × 300 μm |
| Conductivity of tissue ($\sigma$) | 0.13 S/m |
| Air dimension ($L \times W \times H$) | 10 mm × 10 mm × 4 mm |
| Energy error (user-specified) | 1 % |
| Final solution no. of mesh elements | 410,000 |
| Adaptive passes (converged) | 6 |

## 3. Results

### 3.1. μMS of MFB Fibers and FSCV Measurements from the Striatum

In this work the MagPen: Type <u>H</u>orizontal (Type H or MagPen-Type H) activated the MFB fibers (see Fig. 1(a)) and the CFM.WINCS *Harmoni* [11] was used to track the evoked *in vivo* dopamine release from the striatum using FSCV (see Fig. 1(a)-i and Fig. 1(a)-ii). The MagPen driving circuitry and the associated waveforms are demonstrated in the following signal flowchart (see Fig. 1(b)). A function generator (Model no. RIGOL DG1022Z) is used to generate bursts of triangular waveforms. This is the current through the μcoil (***i(t)***) (denoted by solid-black colored lines in Fig. 1(b)-i). Then the current ***i(t)*** is amplified by a Class-D amplifier (Model no. Pyramid PB717X) set at a constant gain A (see Fig. 1(b)-ii). This time-varying current through a μcoil generates a time-varying magnetic field (***B(t)***) (represented by black-dotted lines in Fig. 1(b)). As per Faraday’s Law of Electromagnetic Induction, this time-varying magnetic field induces an electric field (***E(t)***) (represented by solid-red lines in Fig. 1(b)-iii) which is used to activate the MFB fibers. This follows directly from the equation: 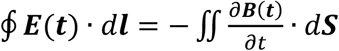. Therefore, we obtain: 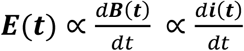. Hence, on applying bursts of triangular current waveform (***i(t)***) through the μcoil, we will obtain an induced electric field waveform (***E(t)***) in the form of bursts of a biphasic square waveform (see solid-red lines in Fig. 1(b)-iii). As this induced electric field activates the MFB fibers, it will induce a biphasic square current, ***J(t)*** (represented by solid green lines) in the MFB fibers, thereby activating them (see Fig. 1(b)-iv). More details on the specifics of the triangular waveform that triggers dopamine release can be found in section 3.3.

Upon stimulation from the μcoils in the MagPen, due to propagation of action potentials along MFB axons, depolarization takes place at post-synaptic striatal dopaminergic neurons. This is followed by the influx of Ca-ions through voltage-gated calcium ion channels (see Fig. 1(c)) triggering exocytosis of dopamine-containing vesicles. The CFM with the assistance of WINCS *Harmoni* tracks this evoked neurotransmitter (in this case, dopamine) release using FSCV (see Fig. 1(c)).

### 3.2. Simulation study of Electric Field for Dopamine Release upon Electric & Micromagnetic Stimulation of the MFB Fibers

In previous studies on evoked dopamine release upon MFB stimulation using electrical stimulation *in vivo*, 2 mA of biphasic square current waveforms of duration 4 ms and frequency ranging between 60-130 Hz were applied through the electrode [30]. The goal was to determine the amplitude of the current through the μcoils that can generate an induced electric field value similar to the electric field generated from electrical stimulation *in vivo*. Using ANSYS-Maxwell (electrostatic solver), we studied the electric field contour plot from the bipolar twisted electrodes (Model no. Plastics One, MS 303/2, Roanoke, VA, USA) upon being driven by 0.2 mA of current (see Fig. 2(a)). The circular wire tips of diameter 0.127 mm separated by 1 mm have been denoted with yellow circles (see Fig. 2(a)). The electric field has been simulated on a tissue of dimension 2 mm × 2 mm × 300 μm. Since the electrodes are in direct electrochemical contact with the tissues during neuromodulation, in this FEM-modeling study in Fig. 2(a) (see Table 1 for the detailed modeling parameters), the distance between the electrical electrode and the biological tissue was negligible.

**Figure 2.**
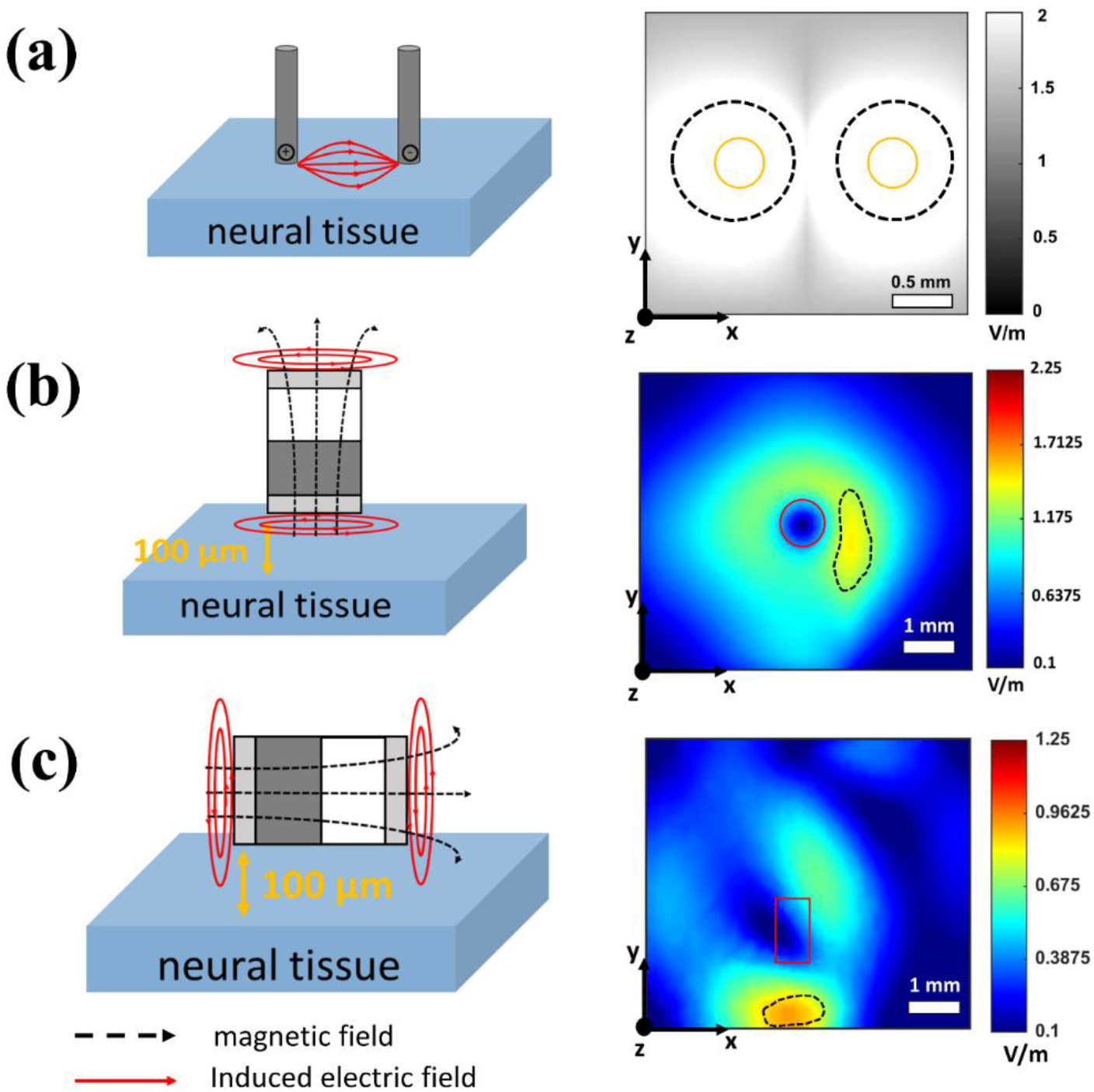
Comparison of the electric field from electrical electrode and magnetic μcoil using finite element modeling (FEM) by ANSYS-Maxwell. (a) Electric field generated on a tissue of dimensions 2 mm × 2 mm × 300 μm from the twisted bipolar electrical electrode (Model no. Plastics One, MS 303/2, Roanoke, VA, USA) with wire tips (shown here with yellow-outlined circles) separated by 1 mm. Current through the electrical electrodes was 0.2 mA. (b) Induced electric field on a tissue of dimensions 4 mm × 4 mm × 300 μm from the MagPen-Type V (shown here with red circle outline). The field has been measured at a distance of 100 μm between the μcoil and tissue. The μcoil is being driven by a sinusoidal current of 5.4 A at 60 Hz frequency. (c) Induced electric field on a tissue of dimensions 4 mm × 4 mm × 300 μm from the MagPen-Type H (shown here with red rectangle outline). The field has been measured at a distance of 100 μm between the μcoil and tissue. The μcoil is being driven by a sinusoidal current of 5.4 A at 60 Hz frequency.

The magnetic μcoils were also numerically simulated on ANSYS-Maxwell (eddy current solver) to match the induced electric field values to that of the electric field values reported in Fig. 2(a). Since the μcoils are not in direct electrochemical contact with the brain tissues (due to the insulating coating), we have simulated a distance of 100 μm between the μcoils and the biological tissue of dimension 2 mm × 2 mm × 300 μm. Fig. 2(b) shows the induced electric field contour plot when the μcoil of the MagPen is oriented in the Type V orientation with respect to the tissues. Fig. 2(c) shows the electric field contour plots when the μcoil of the MagPen is oriented in the MagPen-Type H orientation with respect to the tissues. It was investigated through FEM modeling that a sinusoidal current of amplitude 5.4 A at a frequency of 60 Hz was necessary to drive the μcoil such that it could generate an induced electric field on a biological tissue located 100 μm away, of similar orders of magnitude as that from electrical stimulation (see Fig. 2(c)).

Upon comparing the electric field contour plots from Fig. 2(a) and Fig. 2(b)-(c), we have shown how much more spatially specific the μcoil is compared to the electrical stimulation electrodes. In Fig. 2(a), the spatially uniform field of activation around the electrode tips spans in cylindrical blocks of volume ∼0.24 mm^3^, each. That calculates to 40 % of the tissue volume which has the maximum probability of activation with the threshold electric field being around 1.8 - 2 V/mm (denoted by black-dotted lines in Fig. 2(a)). Whereas in Fig. 2(b) & Fig. 2(c), only 1.13 mm^3^ and 0.63 mm^3^ of the tissue volume out of 4.8 mm^3^ (denoted by black-dotted outlines) has maximum potential of getting activated by MagPen-Type V & MagPen-Type H, respectively. This calculates to a spatial specificity of ∼24% and 13% of the tissue volume for MagPen-Type H and MagPen-Type V prototypes, respectively (see Fig. 2(b) & Fig. 2(c)). If one rotates the μcoil in a clockwise or in an anticlockwise direction keeping the tissue location constant, it will activate a completely different volume of tissue. Since the induced electric field values in the spatial contour plots are greater for MagPen-Type V, it might be a common notion that Type V would successfully activate the MFB fibers and Type H will not. However, our experiments show that Type H successfully activated MFB fibers whereas Type V did not. Herein lies the importance of directionality of the induced electric field in successfully activating the MFB fibers. In section 3.4, we have discussed the clinical significance of this directionality and/or orientation dependence of magnetic μcoils with respect to the neural fibers.

### 3.3. Current Waveform Driving the μcoil in MagPen to Activate the MFB Fibers

Previous reports suggest that a current (***J(t)***, see Fig. 3(a)) of amplitude 300 μA in the shape of a biphasic square of duration 4 ms (= 2 ms duration in each phase) and frequency 60 - 130 Hz needs to be injected into neural tissues to evoke dopamine release *in vivo* when the MFB fibers are activated by an electrical stimulation [17,31,36]. Keeping this as the standard, we tried to deliver appropriate current waveforms through the μcoil (***i(t)***, see Fig. 3(a)) in the MagPen such that we can correlate the evoked dopamine response from micromagnetic stimulation to that of electrical stimulation *in vivo*. The amplitudes of the induced electric field necessary to activate the MFB for potential dopamine release has been numerically calculated in Fig. 2. We have shown how variation of this amplitude of the current through the μcoil influences the evoked dopamine release from the striatum in section 3.5. This section focuses more on the duration and frequency components of the current through the μcoils.

**Figure 3.**
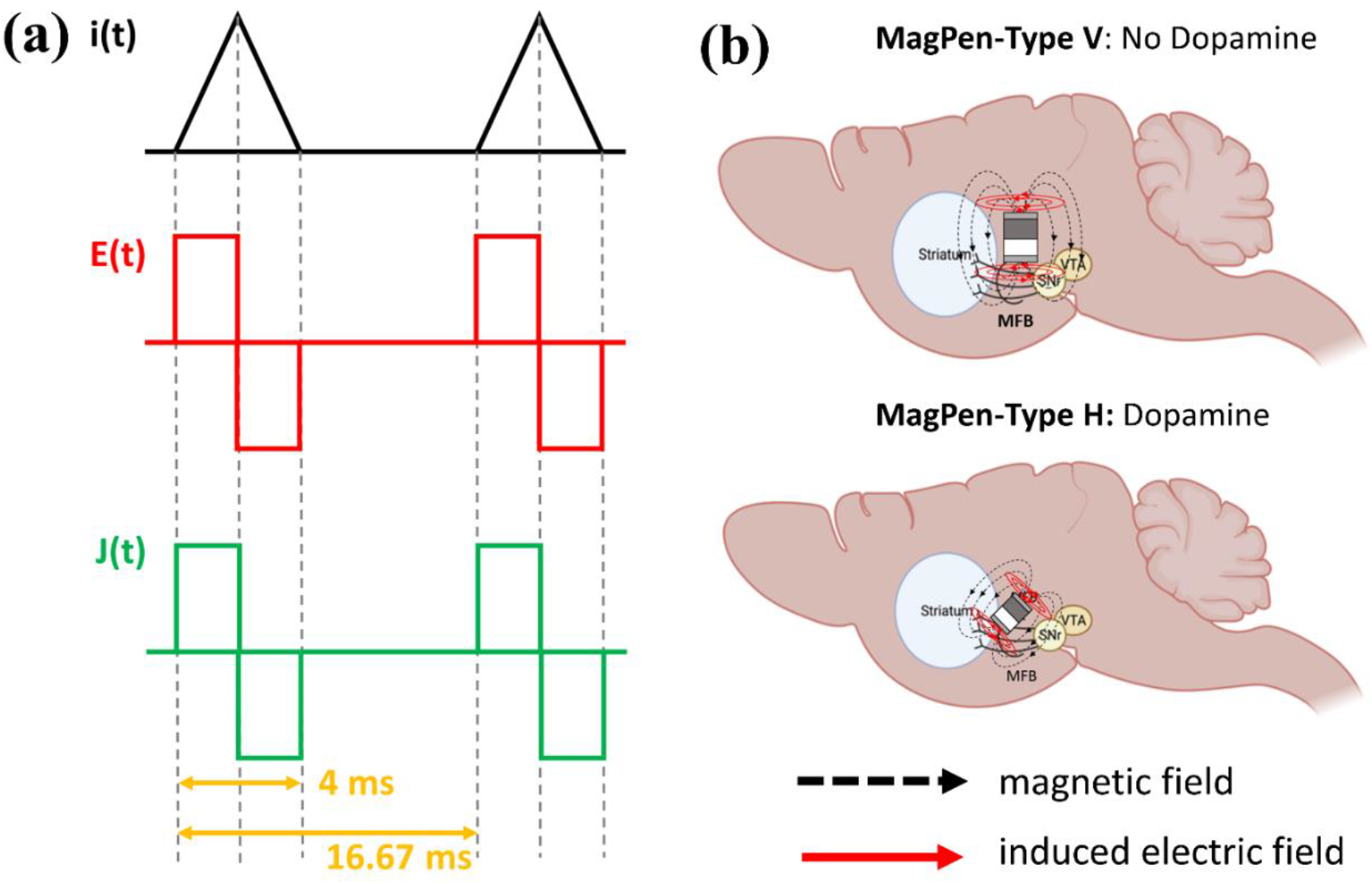
(a) The μcoils in MagPen were driven by bursts of a triangular waveform, ***i(t)*** of duration 4 ms and a frequency of 60 Hz to activate the MFB fibers in the rodent brain. The amplitude was variable. This generated a biphasic induced electric field, ***E(t)*** which in turn induced current, ***J(t)***, in the brain tissues of duration 4 ms and a frequency of 60 Hz. (b) Schematic diagram of MagPen-Type V & Type H oriented over the MFB fibers at the striatum-dopamine pathway. Type V orientation over the MFB did not show any evoked dopamine response, whereas Type H orientation over the MFB (long axis of the μcoil and the MFB fibers perpendicular to each other) showed evoked dopamine response, *in vivo*. Upon stimulation at the MFB, both Type H & Type V showed rat whisker movements. Partially created with BioRender.com.

The μcoils in MagPen being driven by a time-varying current, ***i(t)*** generates a magnetic field, ***B(t)*** which is given by equation (1):

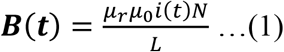

where *μ*_*r*_ is the relative permeability of the medium, *μ*_0_ is the vacuum permeability, *N* is the number of turns of the μcoil and *L* is the length of the μcoil. Fig. S3 of Supplementary Information S3 contains the spatial contour graph for the magnetic flux density (***B***_***x***,***y***,***z***_). However, from equation (1) it is evident that the temporal component of the magnetic field is equivalent to that of the current driving the μcoil.

Following the rules of Faraday’s Law of Electromagnetic Induction, this time-varying current induces an electric field on a conductive medium, which, in this work, is the brain. This induced electric field can be expressed by equation (2):

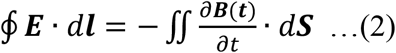

where ***B***(***t***) is the magnetic field generated from the μcoil, ***E*** is the induced electric field activating the neurons, and ***l*** and ***S*** are the contour and the surface area of the neural tissue.

Upon substituting the value of ***B***(***t***) from equation (1) in equation (2), we get the following in equation (3):

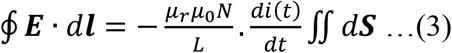

Therefore, we get that the required induced electric field for neural activation from the μcoil is a first order time-derivative of the time-varying current through the μcoil. Recalling our previous statement in this section, we required the current through the neural tissues to be a biphasic square waveform (***J(t)***, represented by solid-green lines in Fig. 3(a)) of duration 4 ms and frequency 60 Hz. This means that the induced electric field also needs to be a biphasic square waveform of the exact same duration and frequency (***E(t)***, represented by solid-red lines in Fig. 3(a)). Since 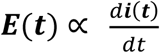, the current (***i(t)***) through the μcoil needs to be a triangular waveform of the same duration and frequency shown in solid-black line in Fig. 3(a). To summarize, we applied 1-cycle bursts of a triangular current waveform of duration 4 ms; each burst was repeated at a frequency of 60 Hz, meaning each burst was separated by a duration of 12.667 ms. The total stimulation applied was for a time span of 2 seconds (trigger controlled by the function generator) and a wait time of 5 min was applied between each successive trial.

### 3.4. Orientation of the μcoil with respect to the MFB Fibers for Evoked Dopamine Release *in vivo*

The correct orientation of the μcoil with respect to neural tissues has been quite a debated topic for the micromagnetic neurostimulation technology [19,20,23,25]. In addition to the spatially specific nature of micromagnetic stimulation (discussed in section 3.1), such directional activation provides a significant advantage of micromagnetic neurostimulation over electrical stimulation. As depicted in Fig. 3(b), MagPen-Type V orientation over the MFB fibers did not evoke dopamine release *in vivo*, whereas Type H orientation over the MFB fibers did. Surprisingly, both Type V and Type H orientations showed rat whisker movements upon *in vivo* stimulation at the MFB. Although the anatomical pathway involved in generating and sensing rat whisker movements is not the primary focus of this work, it was used as a metric to figure out whether the electric field was being applied to the MFB of the rat brain [37].

Fig. 4 provides an explicit explanation for such μcoil-orientation dependent dopamine release from the rat striatum. The way the μcoil is oriented with respect to the MFB fibers play a critical role in determining successful activation of the MFB fibers. The successful activation upon any kind of external stimulation depends on the neuron membrane hyperpolarization and depolarization effect [38]. During electrical stimulation using bipolar electrodes, one electrode terminal being positive and the other being negative causes membrane depolarization and membrane hyperpolarization on the MFB fibers respectively (see Fig. 4(a)-i). Hence, this activates the MFB fibers to release dopamine into the striatum. The phenomenon is better explained with the electric field spatial contour plots and directionality of the electric field in Fig. 4(a)-ii. Similarly, when the long axis of the μcoil is oriented perpendicular to the MFB fibers (this is achieved by MagPen-Type H), the directionality of the induced electric field is like that of the bipolar electrical electrodes (see Fig. 4(b)-i). The MagPen-Type H could also be oriented over the MFB fibers such that the long axis of the μcoil is parallel to the MFB fibers (see Fig. 4(c)-i). However, the directionality of the induced electric field for Fig. 4(c)-i is such that it does not support membrane hyperpolarization and depolarization that could activate the MFB fibers. Therefore, although the contour plots for the induced electric field is same for both Fig. 4(b)-ii and Fig. 4(c)-ii cases, the directionality of the induced electric field for the orientation in Fig. 4(c) does not support MFB activation, whereas that in Fig. 4(b) does. When MagPen-Type V is oriented over the MFB fibers, the induced electric field directionality forms a closed circuit that is unable to activate the MFB fibers (see Fig. 4(d)-i & Fig. 4(d)-ii). This is the reason why we saw evoked dopamine release from the striatum with MagPen-Type H and not with MagPen-Type V stimulation over the MFB fibers.

**Figure 4.**
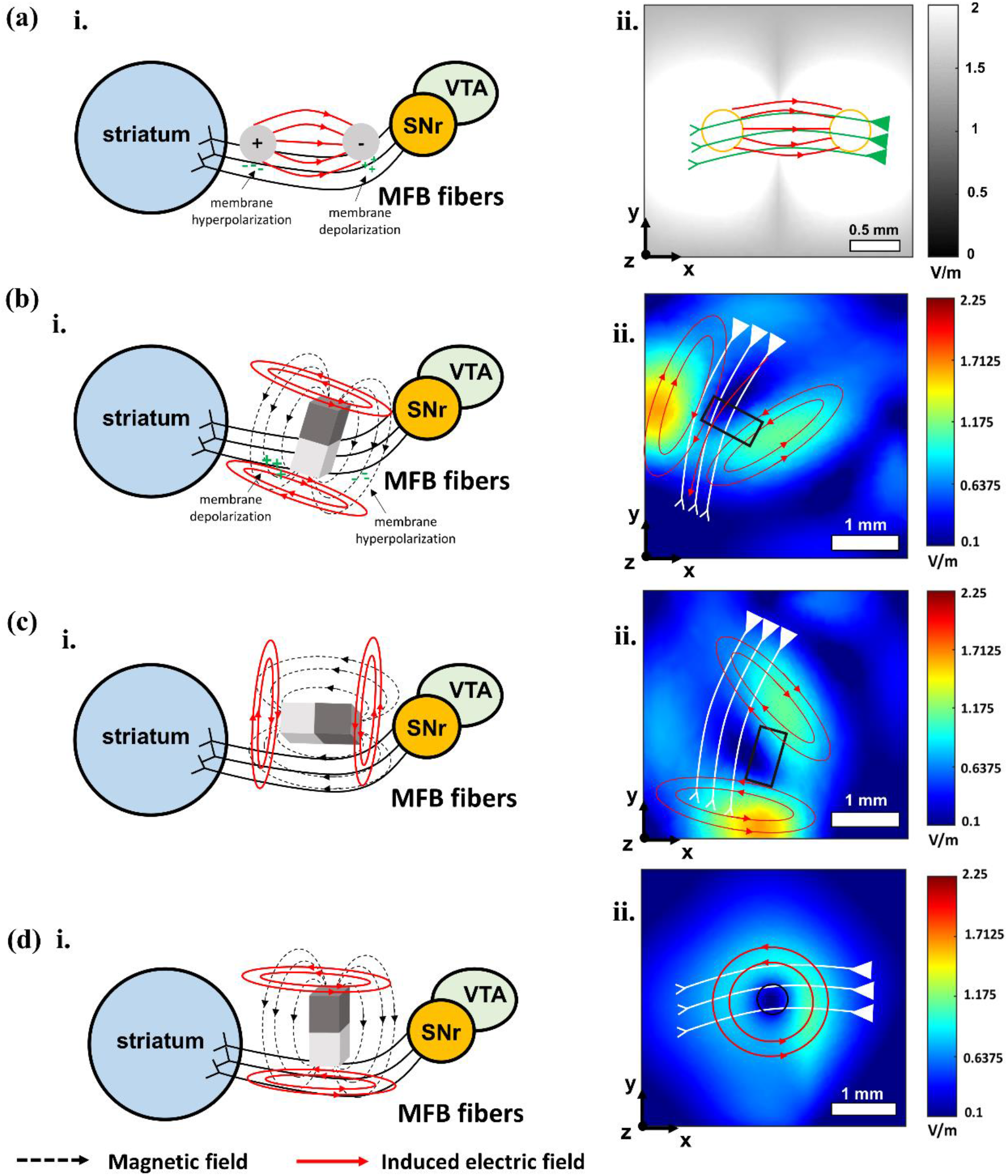
(a) i. Membrane hyperpolarization and depolarization when MFB fibers are stimulated using bipolar twisted electrical stimulation electrodes. ii. Spatial contour plots for electric field from the bipolar electrodes. The outline for the tip of the bipolar electrodes is shown in yellow circles. The MFB fibers are shown in green. (b) i. Membrane hyperpolarization and depolarization when the long axis of the μcoil is oriented perpendicular to the MFB fibers. This is the orientation of MagPen-Type H with respect to the MFB fibers that activated the MFB-striatum dopamine pathway in our *in vivo* experiments. ii. Spatial contour plots for induced electric field from MagPen-Type H. (c) i. No membrane hyperpolarization or depolarization was observed when the long axis of the μcoil was oriented parallel to the MFB fibers (also achieved by MagPen-Type H); hence there was no evoked dopamine release. ii. Spatial contour plots for induced electric field from MagPen-Type H. (d) i. No membrane hyperpolarization or depolarization was observed when MagPen-Type V oriented over the MFB fibers; hence there was no evoked dopamine release. ii. Spatial contour plots for the induced electric field from MagPen-Type V. For (b)-(d) ii. The outline for the μcoil is shown in red. The MFB fibers are shown in white.

### 3.5. Amplitude Dependent Control of Dopamine Release

The MagPen-Type H stimulator was tested *in vivo* in a male Sprague-Dawley anesthetized rat (200g). Briefly, the MagPen was lowered into the right medial forebrain bundle (MFB; coordinates from Bregma in mm: AP -4.6, ML 1.3, DV -8), and the CFM was lowered to the right dorsal striatum (coordinates from Bregma in mm: AP 1.3, ML 2.0, DV -4). Micromagnetic stimulation was applied to the MFB while evoked dopamine release was measured in the dorsal striatum. The stimulation parameters for micromagnetic stimulation were custom calculated to suit the stimulation parameters typically used for FSCV to evoke dopamine release with bipolar electrode stimulation (biphasic square waveform of 300 μA amplitude, 2 ms pulse width and 60 Hz frequency (see Fig. 3(a))).

The current required to drive the μcoil (***i(t)***) and the corresponding current injected into the brain tissue for dopamine release (***J(t)***) is correlated (see Supplementary Information S2). From our calculations, to inject a current (***J(t)***) of 300-500 μA in to the MFB, we would need the μcoil to drive an approximate current (***i(t)***) between 3 and 5.4 A. Stimulation was delivered at progressively increasing potentials, with the value of ***i(t)*** varying from 3 A to 5.4 A (see Fig. 5(a) & (b)). Representative voltammograms (see Fig. 5(a)-i & (b)-i), oxidative current traces (see Fig. 5(a)-ii & (b)-ii), and FSCV pseudocolor plots (see Fig. 5(a)-iii & (b)-iii) after stimulation at 3 A and 5.4 A are shown in Fig. 5(a) & (b). Averaging across 3 different trials, it is seen that increasing stimulation amplitude leads to significantly increased dopamine release in the dorsal striatum (paired t-test, 95 nM to 210 nM, *p = 0*.*031*) (see Fig. 5(c)).

**Figure 5.**
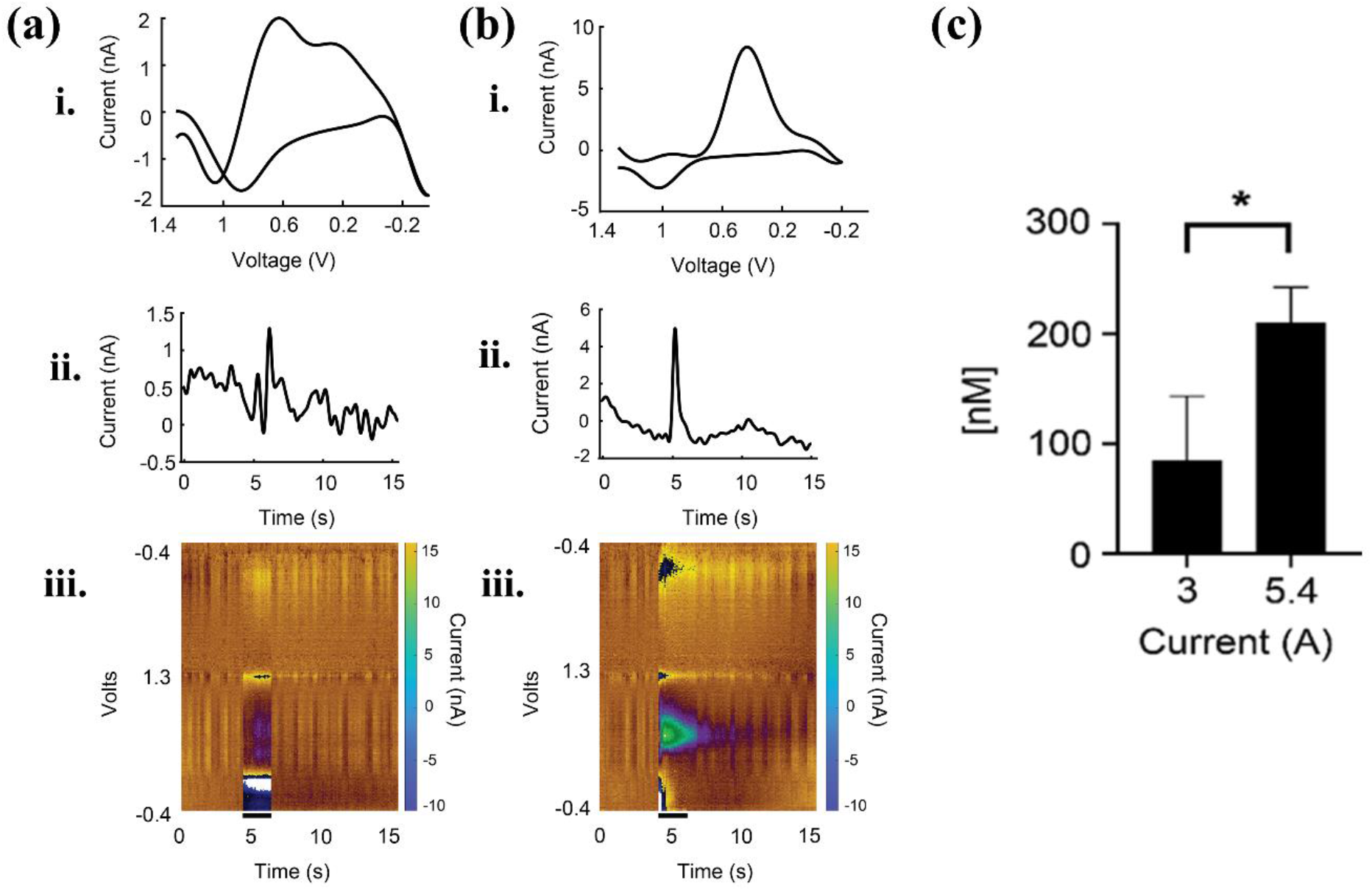
*In vivo* evoked release of dopamine. Micromagnetic stimulation was applied to the MFB via the MagPen stimulator while evoked dopamine release was measured at the striatum. Representative figures of stimulation at (a) ***i(t)*** = 3 A at 4 ms pulse width and 60 Hz frequency (triangular) and (b) ***i(t)*** = 5.4 A at 4 ms pulse width and 60 Hz frequency (triangular). Voltammograms ((a) & (b)-i), oxidative current changes ((a) & (b)-ii), and FSCV pseudocolor plots ((a) & (b)-iii) are shown. (c) Summary data at different stimulation potentials (*n = 3*). Errors are the standard deviations. Black bar represents the duration of magnetic stimulation. *Denotes statistical significance, *p = 0*.*031*.

## 4. Discussion

This is the first report of micromagnetic stimulation of the MFB fibers *in vivo* to study the evoked dopamine release from the striatum. Micromagnetic stimulation technology is a new neuromodulatory intervention and is still in its infancy. A study of the neurotransmitter release associated with this therapeutic intervention is needed. This work is an early stage study of the same where we used the MagPen as the micromagnetic stimulator to activate the MFB fibers. We studied the *in vivo* evoked dopamine release from the striatum using FSCV. Our study confirmed the importance of the directionality of micromagnetic stimulation. Only one orientation where the long axis of the μcoil is oriented perpendicular to the MFB fibers could successfully activate the MFB fibers to induce dopamine release *in vivo* into the striatum. This orientation could be achieved by MagPen-Type H prototype. Here lies the importance of the directionality of the induced electric field from these μcoils.

In this work, we tried to design the μcoil driving waveforms such that they corroborate those of the electrical stimulation waveforms for MFB activation and dopamine release. Therefore, we designed bursts of triangular waveform of duration 4 ms and 60 Hz frequency such that the induced electric field waveform is a biphasic square waveform of the same duration and frequency. By altering the amplitude of the current driving the μcoil, we could successfully control the *in vivo* dopamine concentration released in the striatum. The concentration of dopamine recorded changed from 95 nM to 210 nM in the striatum when the amplitude of the triangular current through the μcoil was changed from 3 A to 5.4 A.

This work being a proof-of-concept study, there is immense scope of improvement which can pave the path for future studies. A major drawback of the MagPen implant was the bulky dimension of the probe. The MagPen-Type H probe was 1.7 mm in width and 0.4 mm in thickness compared to the bipolar electrodes where each wire had a diameter of 0.127 mm separated by 1 mm. So to insert the MagPen-Type H probe in the MFB of rodent brains *in vivo*, a rectangular hole of length 1.7 mm and width 0.4 mm was required to be drilled. Whereas, to insert the bipolar electrodes in the brain only a hole of length 1.3 mm and width < 0.2 mm is usually required to be drilled. Therefore, *in vivo* MagPen insertion in the MFB was an extremely delicate process as it had the possibility to induce rodent brain tissue damage and bleeding. To work around this drawback of the micromagnetic stimulation technology, there is an ardent need for a new design of the μcoil which is fabricated on a flexible material (i.e. there is no density mismatch between the substrate material of the μcoil and the brain tissues). In addition, more efficient μcoils, even μcoil arrays, need to be studied and designed such that they offer more spatially selective activation, thereby improving the focality of micromagnetic neurostimulation even more. Furthermore, as it is unknown if the increase in dopamine concentration with increase in the amplitude of micromagnetic stimulation is linear, this motivates future experiment design in this line of work.

## 5. Conclusions

We have shown that *in vivo* micromagnetic neurostimulation of the MFB using μcoils of dimension 1 mm × 500 μm × 600 μm can trigger dopamine release from the striatum. The WINCS *Harmoni* system working on the principle of FSCV successfully tracked this dopamine release *in vivo* using carbon fiber microelectrodes (CFM) in real time. Only MagPen-Type H could successfully activate the MFB fibers for dopamine release in the striatum, that too when the long axis of the μcoil was oriented perpendicular to the MFB fibers. Therefore, other than the magnitude of the induced electric field that determines successful activation of the fibers, the directionality of the induced electric field from these μcoils is also key to successful activation. Furthermore, increasing the amplitude of micromagnetic neurostimulation by 1.8× alters the concentration of dopamine released in the striatum by 2.2×. Overall, this work proves the efficiacy of micromagnetic stimulation as an alternative to electrical stimulation in activating the dopamine neurochemical pathway.

## Acknowledgements

This study was financially supported by the Minnesota Partnership for Biotechnology and Medical Genomics under award number ML2020. Chap 64. Art I, Sec11on 4. R.S. acknowledge the 3-year College of Science and Engineering (CSE) Fellowship awarded by University of Minnesota, Twin Cities. Portions of this work were conducted in the Minnesota Nano Center (MNC), which is supported by the National Science Foundation through the National Nano Coordinated Infrastructure Network (NNCI) under Award Number ECCS-2025124. J.P.W and R.S. also thank the Robert Hartmann Endowed chair support. Research reported in this publication was supported by the University of Minnesota’s MnDRIVE (Minnesota’s Discovery, Research and Innovation Economy) initiative. This research was also supported by the National Institutes of Health (NIH) R01NS112176 (Y.O., K.L.), R42NS125895-01A1(K.L., K.B.), and R01NS129549 (H.S., Y.O., K.L.) awards.

