## Supplementary Information for "Micromagnetic Stimulation (μMS) Controls Dopamine Release: An *in vivo* Study Using WINCS *Harmoni*"

### S1. The complete Magnetic Pen (MagPen) prototype & RLC measurements of the µcoil

Table S1. RLC measurements of the µcoil

| Parameters | Value @ 1 kHz | Value @ 120 Hz |
|---|---|---|
| Type of coil | solenoid | solenoid |
| No. of turns (N) | 21 | 21 |
| DC Resistance $R_{DC}$ | 1.8 – 2.2 Ω | 1.8 – 2.2 Ω |
| Series Inductance $L_s$ | 0.45 – 0.467 µH | 0.345 – 0.494 µH |
| Series Capacitance $C_s$ | 52.1 – 53.34 mF | 3.4 – 4.8 F |
| Parallel Inductance $L_p$ | 139.1 – 145.5 mH | 10.68 – 14.21 H |
| Parallel Capacitance $C_p$ | 181.2 – 182.5 nF | 109 -127.5 nF |

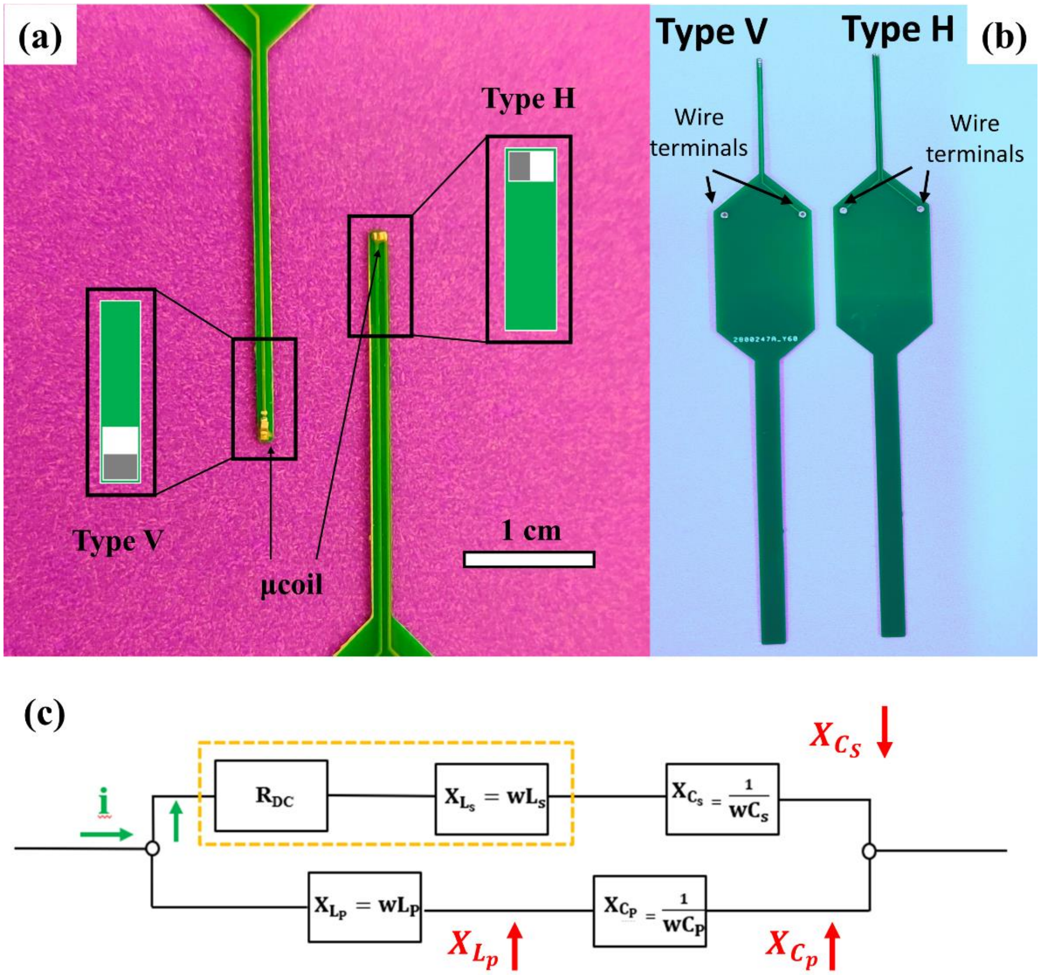


Figure S1. (a) Zoomed-in image for the MagPen: Type Horizontal (Type H) and MagPen: Type Vertical (Type V). The prototypes have been fabricated for implantation in the MFB of rodent brain. The µcoil implant is situated at the tip. (b) Complete prototype for the MagPen, both Type H and Type V, with wire terminals. (c) The circuit equivalent of MagPen reducing to that of a series RL circuit.

Resistance (R), inductance (L) and capacitance (C) measurements of the µcoil (n = 5) at 1 kHz & 120 Hz from LCR meter (Model no. BK Precision 889B), (see Table 1) showed that the electrical circuit equivalent of the µcoil within this frequency range is a series RL circuit. In this work, the frequency range at which the µcoils were being driven to activate the MFB was about 250 Hz. Hence, the LCR meter measurements were carried out at 120 Hz and the other at 1 kHz.

Reactance due to capacitance is given by $\frac{1}{wC}$, while reactance due to inductance is given by $wL$; where $w = 2\pi f$ and $f$ is the frequency at which the measurement is made, here 1 kHz. Therefore, higher the capacitance, lower is the impedance, implying less resistance to the flow of current. But higher the inductance, higher is the impedance, implying greater resistance to the flow of the current.

Ohm's Law, $V = iR$ , where V = voltage, i = current and R = resistance, states that current always travel through the path of lowest resistance in an electrical circuit. From the RLC parameters of the µcoil measured in Table 1 (the measurements varied with an error margin of ±0.0001% between different µcoils due to difference in soldering conditions), the series capacitance ($C_s$) being small (for both frequency measurements) offer low resistance to the flow of the current (see Fig. S1(c)). While the parallel inductance ($L_p$) being high and the parallel capacitance ($C_p$) being low (for both frequency measurements), both offered higher resistance to the flow of current (see Fig. S1(c)). Therefore, whatever current was applied to the µcoil travelled through the path of lowest impedance, the resistance, and the series inductance ($L_s$) (see Fig. S1(c)).

**S2. Correlation between the current driving the µcoil (i(t)) and the current injected into the brain tissues (J(t)) – An estimate**

In µMS, we estimate the neuron activating capability of the µcoil using the induced electric field from the µcoil (***E(t)***). This ***E(t)*** is directly correlated to the time-varying current driving the µcoil (***i(t)***). Therefore, to correlate µMS to that of electrical stimulation to activate the dopamine neurochemical pathway, we required to understand the electric field magnitude from the bipolar electrodes that activate the MFB fibers.

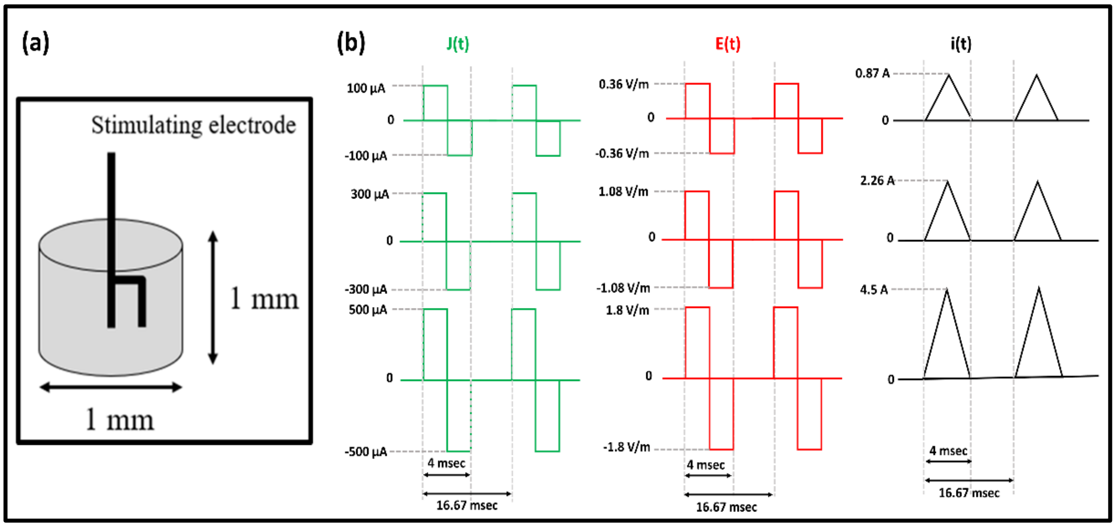


Figure S2.(a) Pictorial diagram of the bipolar electrode activating a cylindrical neural tissue area of height 1 mm and diameter 1 mm. (b) Calculation of the current driving the µcoil (***i(t)***) from 3 different values of the current injected into the neural tissue (***J(t)***) (100 µA, 300 µA and 500 µA) through electric field induced in a cylindrical shaped neural tissue of height 1 mm and diameter 1 mm.

As pictorially demonstrated in Fig. S2(a), we considered a cylindrical tissue area of 1 mm diameter and 1 mm height to be activated. The resistance of the neural tissues can be averaged as R = 3.6 Ω [1–3]. For a current of 300 µA injected into the neural tissues (=***J(t)***) to observe dopamine releases from the striatum [4], the voltage applied to the neural tissues can be calculated to be, V = iR = (300 µA) × 3.6 Ω = 0.00108 V. Therefore, to activate a tissue area of length $\Delta d = 1\,mm$, the electric field necessary can be estimated as: $E = \Delta V/\Delta d$. Hence, ***E(t)*** = 1.08 V/m. Therefore, to obtain an induced electric field (***E(t)***) value of 1.08 V/m of a specific duration and pulse width we can design the current driving the µcoil (***i(t)***), it's amplitude, duration and frequency, accordingly. Therefore, our back-calculation for the induced electric field estimate follows the path: $\boldsymbol{J(t)} \to \boldsymbol{E(t)} \to \boldsymbol{i(t)}$. Fig. S2(b) pictorially demonstrates the variation of the induced electric field values (E(t)), hence the current driving the μcoil (i(t)), for 3 different values of the current injected through the neural tissue (J(t)).

### S3. Spatial components of the magnetic flux density for MagPen

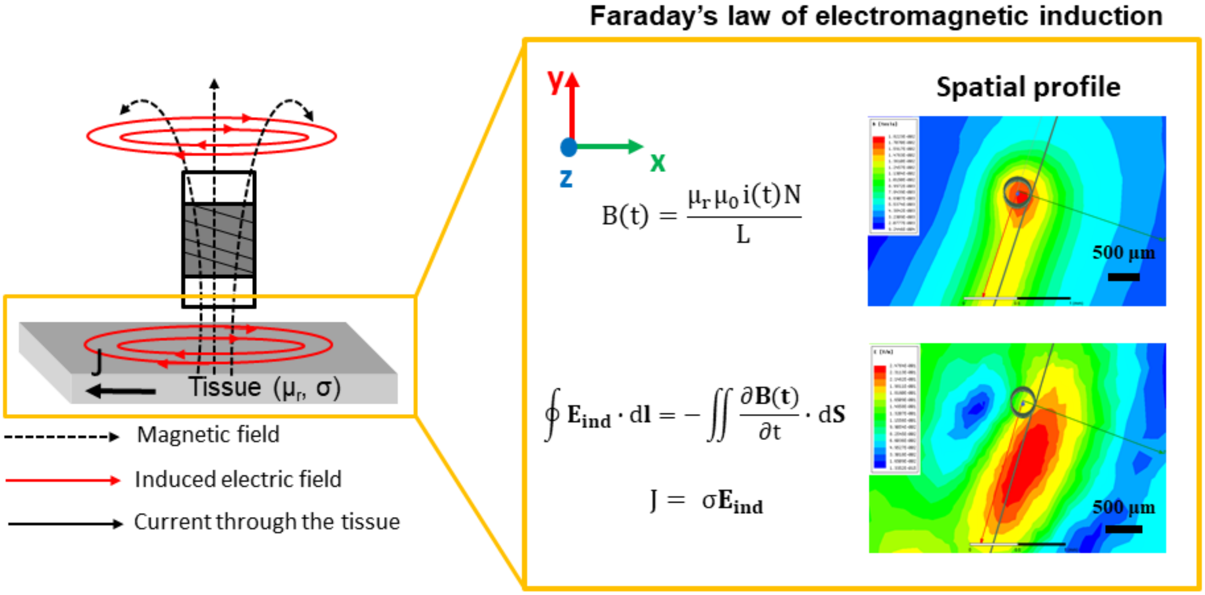


Figure S6. The spatial components of magnetic flux density, ***B(t)*** and induced electric field ($\mathbf{E}_{ind}$) measured for the µcoil and neural tissue distance 300 µm for a sinusoidal current, ***i(t)*** of amplitude 2 A and frequency 2 kHz. Modeling performed on ANSYS-Maxwell (eddy current solver). $\mu_r$ and $\mu_0$ are the relative permeability of the neural tissue and vacuum permeability respectively. σ is the conductivity of the neural tissue. The current that will flow through a neural tissue on stimulation by the induced electric field from these µcoils will depend on σ. When current flows through the neural tissue, the neurons start firing, i.e., it is stimulated.
